# Cytoplasmic polyadenylation by TENT5A is required for proper bone formation

**DOI:** 10.1101/2020.08.18.256115

**Authors:** Olga Gewartowska, Goretti Aranaz Novaliches, Paweł S Krawczyk, Seweryn Mroczek, Monika Kusio-Kobiałka, Bartosz Tarkowski, Frantisek Spoutil, Oldrich Benada, Olga Kofroňová, Piotr Szwedziak, Dominik Cysewski, Jakub Gruchota, Marcin Szpila, Aleksander Chlebowski, Radislav Sedlacek, Jan Prochazka, Andrzej Dziembowski

## Abstract

Osteoblasts orchestrate bone formation by secreting dense, highly cross-linked type I collagen and other proteins involved in osteogenesis. Mutations in Col1α1, Col1α2, or collagen biogenesis factors lead to the human genetic disease, osteogenesis imperfecta (OI). Herein, we show that the TENT5A gene, whose mutation is responsible for poorly characterized type XVIII OI, encodes an active cytoplasmic poly(A) polymerase regulating osteogenesis. TENT5A is induced during osteoblast differentiation and TENT5A KO osteoblasts are defective in mineralization. The TENT5A KO mouse recapitulates OI disease symptoms such as bone fragility and hypomineralization. Direct RNA sequencing revealed that TENT5A polyadenylates and increases expression of Col1α1 and Col1α2 RNAs, as well as those of other genes mutated in OI, resulting in lower production and improper folding of collagen chains. Thus, we have identified the specific pathomechanism of XVIII OI and report for the first time a biologically relevant post-transcriptional regulator of collagen production. We further postulate that TENT5A, possibly together with its paralogue TENT5C, is responsible for the wave of cytoplasmic polyadenylation of mRNAs encoding secreted proteins occurring during bone mineralization.

## Introduction

Bone formation or osteogenesis is a very complex process in which osteoblasts play a crucial role. The primary function of these cells of mesenchymal origin is the secretion of non-mineralized bone matrix (osteoid), whose predominant component is collagen type I (Bilezikian et al., 2008). The collagen fibers form a scaffold on which, with the help of proteoglycans, hydroxyapatite crystals mineralize. Collagens are also secreted by many other types of cells, which makes these fiber-forming proteins the most abundant proteinous constituent of the human body (Brinckmann, 2005). Importantly, mutations affecting collagen I production lead to the human disease osteogenesis imperfecta (OI) (Chu et al., 1983), which comprises a phenotypically and biochemically heterogeneous group of heritable disorders of connective tissue. Characteristic features of OI are skeletal abnormalities leading to bone fragility and frequent fractures, and in some cases short stature and deformations. (Besio et al., 2019; Rauch and Glorieux, 2004). Mutations in *COL1A1* or *COL1A2*, resulting in a quantitative or qualitative defect in type I collagen formation, are responsible for approximately 90% of all OI cases (Van Dijk and Sillence, 2014). Recent studies have led to the discovery of many new, non-collagenous genes causative of OI, most of which are required for collagen I synthesis (Besio et al., 2019), as well as genes involved in bone mineralization and osteoblast homeostasis.

Collagen I undergoes complex posttranslational processing in the endoplasmic reticulum and the Golgi apparatus, which includes hydroxylation, glycosylation, and formation of a triple helix composed of two Col α1(I) chains and one Col α2(I). After secretion, the ends of pro-collagen polypeptides are processed by dedicated proteases, and finally, long collagen fibers are formed. These processing steps are relatively well understood (Bilezikian et al., 2008)

In contrast to the posttranslational phase of α1(I) and α2(I) chain biogenesis, very little is known about the regulation of collagen expression at the mRNA level, although some proteins presumably involved in the stabilization of collagen mRNA, such as LARP6, have been identified (Cai et al., 2010; Zhang and Stefanovic, 2016). However, LARP6 KO mice do not show prominent defects in bone formation (Dickinson et al., 2016). Recently, several patients with OI carrying a mutation in the TENT5A gene were identified (Doyard et al., 2018). TENT5A is a paralogue of the cytoplasmic poly(A) polymerase TENT5C, which acts as an onco-suppressor in multiple myeloma (Mroczek et al., 2017). Importantly, TENT5C increases the expression of immunoglobulins and other secreted proteins in the B cell lineage by stabilizing their mRNAs (Bilska et al., 2020; Mroczek et al., 2017). This may suggest that, by analogy, TENT5A regulates via the same mechanism the expression of secreted proteins in osteoblasts, which dysfunction could lead to a bone-related phenotype.

Here, we show that TENT5A is indeed a cytoplasmic poly(A) polymerase expressed in osteoblasts and osteocytes. Comprehensive analysis of TENT5A KO mouse revealed severe skeletal abnormalities, including frequent bone fractures appearing during or shortly after birth, hypomineralization, short posture, and deformations. Importantly, analysis of the global poly(A) tail distribution by nanopore direct RNA sequencing (DRS) of primary osteoblast cultures revealed that TENT5A polyadenylates and enhances the mRNA expression of both Col1α1 and Col1α2, but also that of other proteins involved in bone formation. In the absence of TENT5A, the production of collagens was drastically decreased. The defect was not only quantitative but also qualitative, as revealed by the aberrant structure of collagen fibers, which correlated with concomitant downregulation of collagen processing enzymes in the TENT5A KO. Therefore, for the first time, we provide a molecular basis for the pathogenesis of TENT5A-related OI (Type XVIII). Additionally, we show that poly(A) tail distribution in osteoblasts undergoes a global change during mineralization, suggesting existence of a wave of cytoplasmic polyadenylation, indispensable for proper bone formation.

## Results

### 1. TENT5A KO mice exhibit a bone-related phenotype with frequent bone fractures

To study the role of TENT5A at the organismal level, we generated TENT5A KO mouse lines in two different backgrounds, inbred C57BL/6N and mixed C57BL/6JxCBA, because in case of TENT5C, a paralogue of TENT5A, genetic background is known to strongly affect the viability of KO mice (Dickinson et al., 2016; Mroczek et al., 2017; Zheng et al., 2019). The CRISPR/Cas9 method was used to establish two mouse lines harboring mutations at the beginning of the second exon (c.436_462del26 in C57BL/6N and c.403_406del4ins50bp in C57BL/6JxCBA), resulting in frameshifts that destroy the catalytic center of the protein (Fig 1A). Initial phenotyping, including microCT analysis, revealed no significant differences between the two strains.

**Figure 1.**
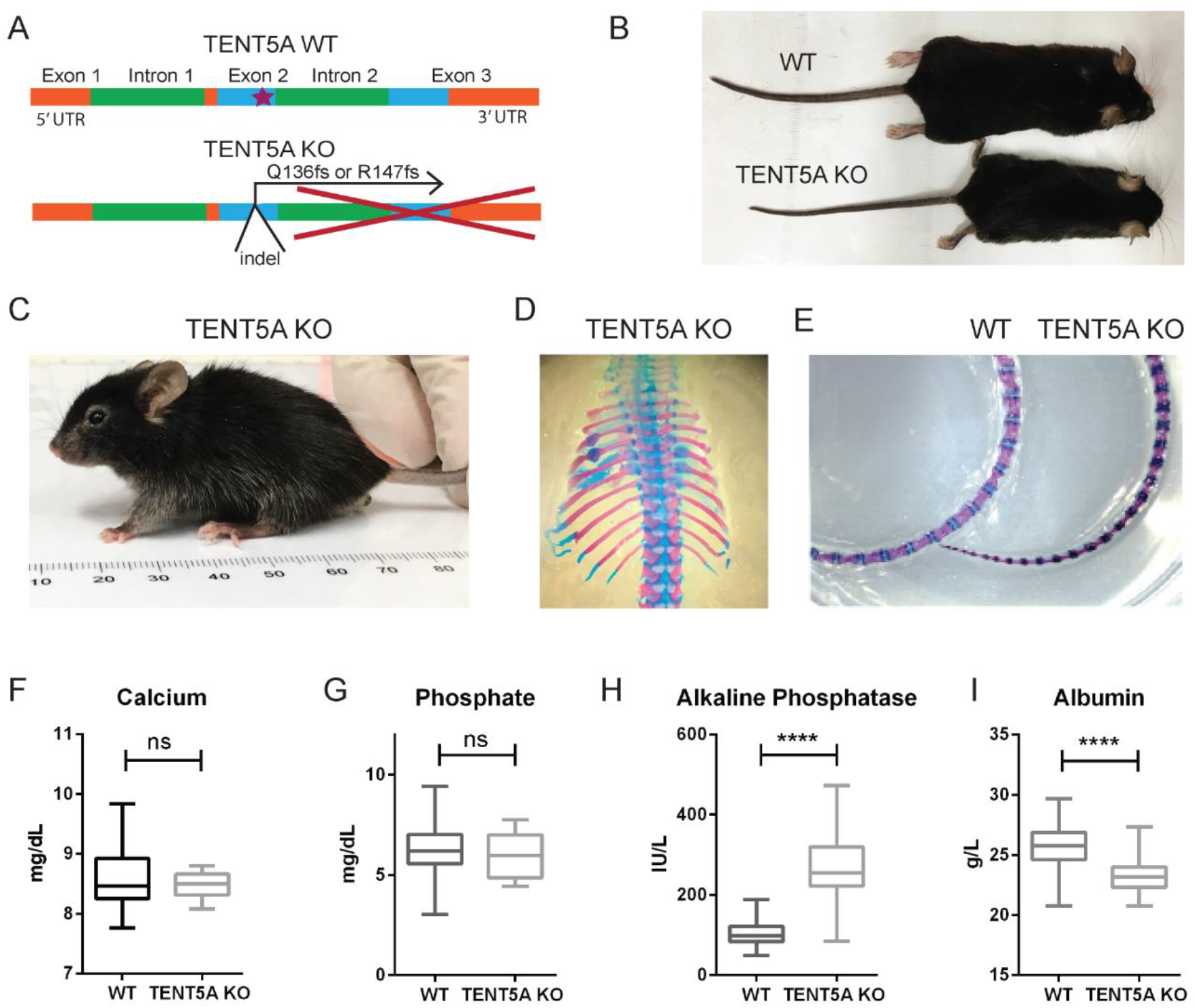
TENT5A KO display phenotypes resembling type XVIII osteogenesis imperfecta. A. Schema of the frameshift mutation introduced by the CRISPR/Cas9 method for the generation of TENT5A KO mice. B. Comparison of the size of adult WT and TENT5A KO littermates. C. Abnormal posture of the TENT5A KO mouse with visible kyphosis. D. Alizarin Red/Alcian Blue staining of a 6-day-old TENT5A KO thorax showing multiple healing bone fractures. E. Alizarin Red/Alcian Blue staining of an 8-week-old WT and TENT5A KO tails reveals decreased cartilage ossification in the TENT5A KO mouse. F–I: Biochemical parameters of TENT5A KO and WT mice. Measurements of calcium (F), phosphate (G), alkaline phosphatase (H) (p<0,0001) and albumin (I) (p<0,0001) in mouse serum (KO, n=15; WT, n=140 – data gathered for the IMPC mouse phenotyping pipeline); Mann–Whitney test.

TENT5A KO animals had slightly decreased survival, which was probably due to perinatal or embryonic lethality, because decreased survival was already visible at P6 and did not change significantly thereafter (Fig EV1A-B). The TENT5A KO mouse was smaller than the WT mouse (Fig 1B, EV1D) and had an abnormal posture with frequent kyphosis and wavy tail (Fig 1C, EV1C). This skeletal phenotype corroborates what was observed previously in mice with the TENT5A mutation generated by ENU-mutagenesis (Diener et al., 2016). However, we also observed frequent bone fractures (Fig 1D, EV1E), which is reminiscent of one of the symptoms of human OI patients harboring the TENT5A mutation (Doyard et al., 2018).

Alizarin Red/Alcian Blue staining of the skeletons of 8-week-old mice revealed multiple rib fractures (Fig EV1E). Occasionally, we detected long bone fractures (Fig EV1F). To determine when fractures first occurred, consecutive staining of newborn mice (P6) and late embryos (E17) was performed. Rib fractures were present in newborn mice (Fig 1D), but not in late embryos (Fig EV1H), suggesting that they appear during or shortly after parturition. Additionally, in adult TENT5A KO mice, we detected decreased cartilage ossification of the tail (Fig 1E).

Biochemical analyses of the TENT5A KO mice revealed that serum calcium and phosphate levels were within normal ranges (Fig 1F-G), as observed in OI type XVIII patients (Doyard et al., 2018). Alkaline phosphatase (ALP) was 2.5-fold higher than in WT mice (Fig 1H). Up until now, the only OI type with an abnormally high ALP level was type VI, which is caused by loss-of-function mutation in the *SERPINF1* gene (Glorieux et al., 2002; Homan et al., 2011). TENT5A KO mice also had low plasma albumin levels (Fig 1I), which was probably a consequence of inflammatory responses to frequent bone fractures.

We concluded that TENT5A KO mice display a bone phenotype typical of OI.

### 2. TENT5A mice exhibit skeletal bone hypomineralization and altered long-bone cortical and trabecular bone microarchitecture

Whole-body scans using micro-computed tomography (μCT) showed multiple morphological abnormalities in the bones of 13–14-week-old TENT5A KO mice and strong hypomineralization over the entire skeleton, as reported previously for the ENU-generated TENT5A mutant (Fig 2A, EV Movie 1-2). In addition, several bone fractures were observed, as well as healed rib fractures, in all of the samples analyzed (Fig 2B). The shape of the ribcage was altered and compressed. Fractures in long bones were less frequent, although broken femurs, the hardest long bone in the body, were sometimes observed (Fig 2C). Notably, the decreased of mineral content is most visible by X-ray absorption heatmap in the paws of TENT5A KO mice, where the blue color stains for hypomineralized loci (Fig 2D).

**Figure 2.**
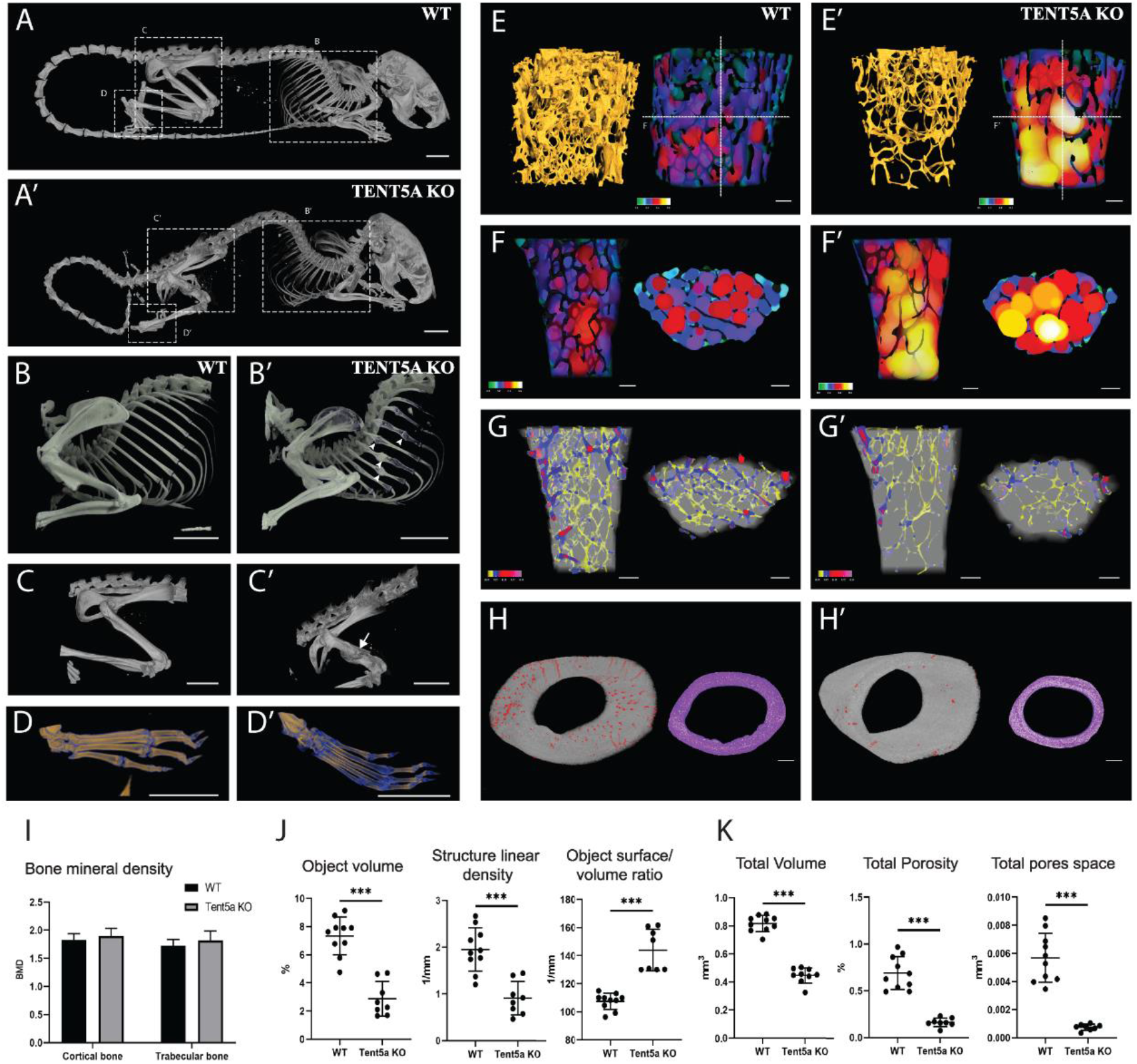
MicroCT analysis reveals skeletal abnormalities of TENT5A KO mice A-H. 3D-MicroCT images of representative WT (A-H) and TENT5A KO mice (A’-H’). A. Whole body hypomineralized skeleton with fractures (dashed squares) and malformations in TENT5A KO mice. Scale bar, 5 mm. B–D. Closer view of several healing ribs fractures (arrowheads) (B, B’). Femur fracture is indicated with the arrow (C, C’) and hypomineralization of paws is displayed in blue (D, D’). Scale bar, 5 mm. E–H. High resolution 3D-MicroCT images of the altered microstructure of distal femoral metaphyseal trabecular bone and mid-diaphyseal cortical bone. Scale bar, 250 μm. E–F. 3D structure of trabecular bone trabeculae showing a bone volume decrease in TENT5A mice is shown in yellow (E, E’). Increase of space between the trabecular bone of TENT5A mice (E, E’, F, F’) is shown in green for the smallest space and yellow-white for the biggest space. Dashed lines represent the sections selected for figure F. G. Longitudinal and transversal sections showing no significant difference in trabecular thickness in shown in color in WT and TENT5A KO mice. H. TENT5A KO Mid-diaphyseal cortical bone showing a decrease in porosity of the bone (left image, pores in red) and altered morphology observed in transversal sections in purple. I-K. MicroCT-derived femur morphometry results. I. Bone mineral density (BMD) analysis of cortical and trabecular bone revealed no significant difference in density. Data is presented as mean ± SD. *P≤0.05, (Two-way Anova test). J. Trabecular bone histomorphometry analysis showing trabecular bone percent object volume (%) and Structure linear density (1/mm) decrease in TENT5A KO mice (1.9521 1/mm± 0.4428 vs. 0.9118 1/mm± 0.3350) and Object surface/volume ratio (1/mm) increase (107.43 1/mm± 5.551 vs. 143.85 1/mm± 13.876). Data are presented as mean ± SD. *P≤0.05, ***P<0.001 (Student’s t-test). K. Cortical bone quantification of total volume of object of interest (mm^3^), total porosity of cortical pores (%), and total volume of pore space (mm^3^) were decreased in TENT5A KO mice. Data are presented as mean ± SD. *P≤0.05, ***P<0.001 (Student’s t-test). Data information: All color bars are displayed as 0 to 1 ratios of the parameter values. Femurs of five WT mice and five TENT5A KO mice were used for the experiment.

High resolution μCT analysis confined to the segmented bone tissue in the femur revealed no significant differences in the bone mineral density of trabecular and cortical bone between TENT5A KO mice and wild-type mice, despite the fact that the whole-body skeleton was hypomineralized (Fig 2I). This was clear indication for possible structural abnormalities in bones rather than mineral composition. To analyze further the structural properties of trabecular and cortical bone (Fig 2E-K, EV2A), we performed volumetric analysis of trabecular bone, which confirmed that the total volume of the trabecular region was not significantly different between WT and TENT5A KO mice (Fig EV2B); however, trabecular bone mass was significantly reduced (Fig 2E, G, EV Movie 3-4). The ratio of trabecular bone volume to the volume of the trabecular region represented as a percentage of the object volume was more than two-fold lower in TENT5A KO mice than in WT mice (7.3451% WT vs. 2.8768% TENT5A, p < 0.05) (Fig 2J). The reduced volume of trabeculae was associated with a decrease in their density and thickness, which resulted in a significantly higher trabecular space in TENT5A KO, strongly affecting the mechanical properties of the bone (Fig 2E, F, EV Movie 3-4).

The second important area responsible for bone mechanic resistance is the bone cortical region. Gross morphology analysis revealed strong shape alteration in TENT5A KO mice, which was best visible in the transverse tomographic section. Unlike the characteristic elliptic shape of WT femurs, TENT5A KO femurs had a circular shape. The total volume of the cortical bone was also 44% lower than in WT mice (0.8161 mm3± 0.0535 in WT vs. 0.4465 mm3± 0.0519 in TENT5A KO, p < 0.05) (Fig. 2H). However, the thickness of the cortical bone itself was not significantly different (Fig EV2B). Moreover, the most striking difference was the strong reduction in the percentage of cortical pores (0.6% in WT vs. 0.2% in TENT5A KO) and their volume (0.0057 mm3± 0.0016 in WT vs. 0.0010 mm3± 0.0009 in TENT5A KO) within the entire cortical bone (Fig 2K).

Bone detailed analysis clearly demonstrated that ultrastructural changes in bones are responsible for general hypomineralized character of TENT5A KO whole skeleton and moreover combination with pore number reduction in cortical bone, which are important for mechanical resistance of bone and causative for frequent bone fractures found in TENT5A KO mice.

### 3. TENT5A KO mineralization defect is recapitulated *in vitro*

The observation that bone mineralization *in vivo* was drastically different in TENT5A KO mice prompted us to examine TENT5A expression in bone tissue. However, despite our best efforts, we could not buy or raise specific antibodies against TENT5A protein. Thus, we generated a TENT5A--3xFLAG mouse line. The animals did not display any detectable phenotypes. We observed TENT5A-3xFLAG expression in osteoblasts and osteocytes (Fig 3A), in agreement with previously reported transcriptomic data (Youlten et al., 2020) and confirmed TENT5A expression in *in vitro* cultured primary osteoblasts derived from neonatal calvaria using two independent approaches: ICC and WB analysis (Figure 3B-C).

**Figure 3.**
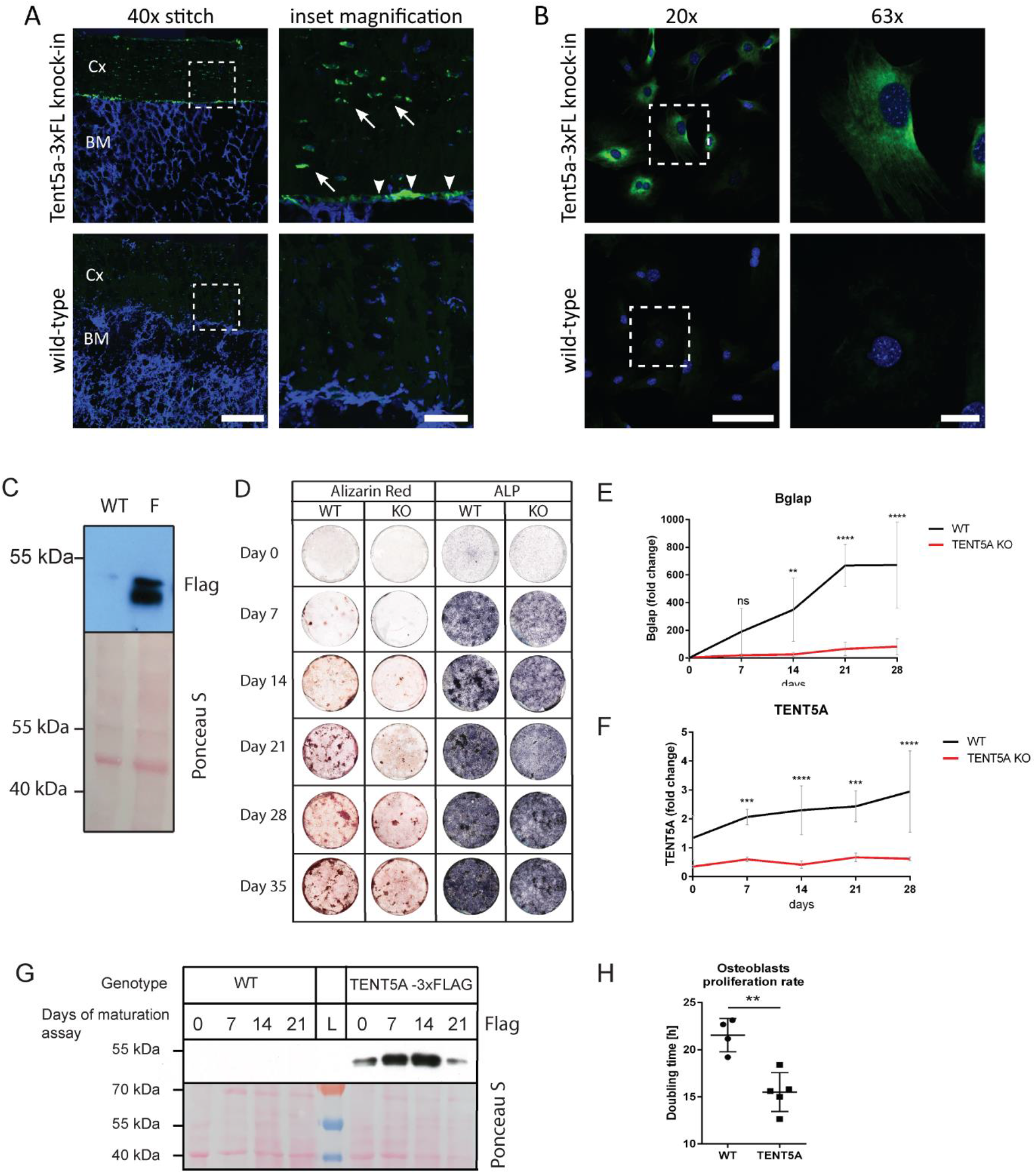
TENT5A is expressed in osteoblasts and regulates the mineralization process A. Immunohistochemical staining for FLAG in femur bone sections from TENT5A-3xFLAG knock-in and wild-type controls with FLAG in green and Hoechst in blue; Cx - cortical bone; BM - bone marrow; arrows and arrowheads indicate FLAG-positive osteocytes within the bone mass and osteoblasts on the bone/bone marrow interface, respectively; scale bars denote 200μm and 50μm on 40x stitch fragment and inset magnification, respectively. B-C: TENT5A-3xFLAG is detectable in TENT5A-3xFLAG, but not in WT osteoblasts. B. Immunofluorescent staining for FLAG in in vitro cultured osteoblasts from Tent5a-3xFLAG and wild-type controls with FLAG in green and Hoechst in blue; scale bars denote 100μm and 20μm for 20x and 63x objective-collected images, respectively. C. Cell lysates from TENT5A-3xFLAG and WT osteoblast neonatal primary cultures were probed with anti-FLAG antibody. Ponceau staining was used as a loading control. D. Osteoblast maturation assay performed on WT and TENT5A KO neonatal calvarial osteoblasts reveals aberrant mineralization in TENT5A KO mice. Cells were stained with Alizarin Red and NBT/BCIP solution on days 0, 7, 14, 21, 28 and 35. E. RT-qPCR analysis of Bglap mRNA levels during osteoblast maturation reveals low levels of Bglap in TENT5A KO mice. Expression was normalized to that of HMBS (n=4–6, medium values from technical triplicates). (Two-way Anova and Sidak’s multiple comparison tests; p-values: D0, ns; D7, ns; D14, 0.0012; D21, 0.0001; D28, 0.0001.) F. RT-qPCR analysis of mRNA levels of TENT5A during osteoblast maturation reveals upregulation of TENT5A expression in the WT, but not in TENT5A KO mouse. TENT5A expression was normalized to that of HMBS (n=4–6, medium values from technical triplicates). (Two-way Anova and Sidak’s multiple comparisons tests; p-values: D0, ns; D7, 0.0008; D14, <0.0001; D21, 0.0001; D28, <0.0001.) G. Western blot analysis of WT and TENT5A-3xFLAG osteoblasts collected on days 0, 7, 14, 21 and 28 of maturation showing upregulation of TENT5A during osteoblast mineralization. H. Analysis of the proliferation rate of adult long-bone derived osteoblasts. Osteoblasts from WT and TENT5A KO mice were stained with CFSE and measured by flow cytometry at 0, 48, and 96 h of culture. Dots represent the average doubling time for each population (n=5–6, p = 0.0022; Unpaired t-test with Welch’s correction).

To examine the mineralization defect in *in vitro* cultures, we established primary calvarial osteoblast cultures from TENT5A WT and KO neonates and performed the maturation assay. NBT/BCIP staining of alkaline phosphatase and evaluation of matrix mineralization by Alizarin Red staining every 7 days showed that osteoblast mineralization was abnormal in TENT5A KO mice (Fig 3D). Mineralized bone nodules were barely present on day 21, and even by day 35, mineralization did not reach the expected level. To confirm compromised differentiation of osteoblasts, we checked the level of *Bglap* (osteocalcin), a marker of mature osteoblasts. Indeed, by day 28 of mineralization, we observed a 670-fold increase in the *Bglap* mRNA level in WT osteoblasts and only an 80-fold increase in KO osteoblasts (Figure 3E).

To assess whether defective osteoblast differentiation and mineralization was caused by loss of TENT5A expression, we examined the level of TENT5A mRNA at different stages in the *in vitro* maturation assay, in TENT5A KO and WT cultures. As expected, the *TENT5A* mRNA level in TENT5A KO osteoblast was residual and did not change over time, whereas the TENT5A mRNA level in WT osteoblasts was higher and increased during osteoblast differentiation (Figure 3F). The marked increase in TENT5A expression during osteoblast differentiation was also visible at the protein level, as visualized in cultures from TENT5A-3xFLAG animals (Figure 3G). Finally, an examination of proliferation rates of osteoblasts isolated from adult calvaria showed that the doubling time of TENT5A KO osteoblasts was significantly lower than that of WT osteoblasts (Figure 3H).

We concluded that TENT5A plays a direct role in osteoblast differentiation and mineralization.

### 4. TENT5A polyadenylates collagen I and other OI-related gene transcripts

Having established that osteoblast cultures recapitulate mineralization defects in TENT5A KO mice, we used this model to dissect further the role of TENT5A in bone mineralization. Initially, we confirmed that, similar to other TENT5 protein family members, TENT5A is an active poly(A) polymerase by using a standard tethering assay (Fig EV3A-B). To determine which mRNAs are regulated by TENT5A polyadenylation, we established primary murine osteoblast cultures and performed genome-wide poly(A) tail profiling using nanopore-based direct full-length RNA sequencing (DRS), which we previously used successfully to identify substrates of TENT5C poly(A) polymerase in B cells (Bilska et al., 2020).

Neonatal WT and TENT5A KO calvarial osteoblasts on D0 and D14 of the maturation assay, in duplicate, were used to generate more than 9 mln mappable transcriptome-wide full-length native-strand mRNA reads (EV Table 1, EV Table 2). We observed no global changes in mRNA polyadenylation status between WT and TENT5A-deficient cells at D0 (Fig 4A), but we did observe subtle shortening of poly(A) tails in TENT5A KO osteoblasts at D14 (Fig 4B). As expected, the majority of the mRNAs encoding housekeeping genes such as components of the translational apparatus or mitochondrial proteins were not affected (Fig EV3C-D). Because TENT5A was upregulated during osteoblast maturation (Fig 3F-G), we examined differences in poly(A) tails between WT and TENT5A KO osteoblasts on D14, which revealed that 52 mRNAs had statistically shorter tails in the TENT5A KO osteoblasts (EV Table 1). Strikingly, genes in which mutations lead to OI and/or are involved in osteoblast differentiation and mineralization were at the top of the list (Fig 4C).

**Figure 4.**
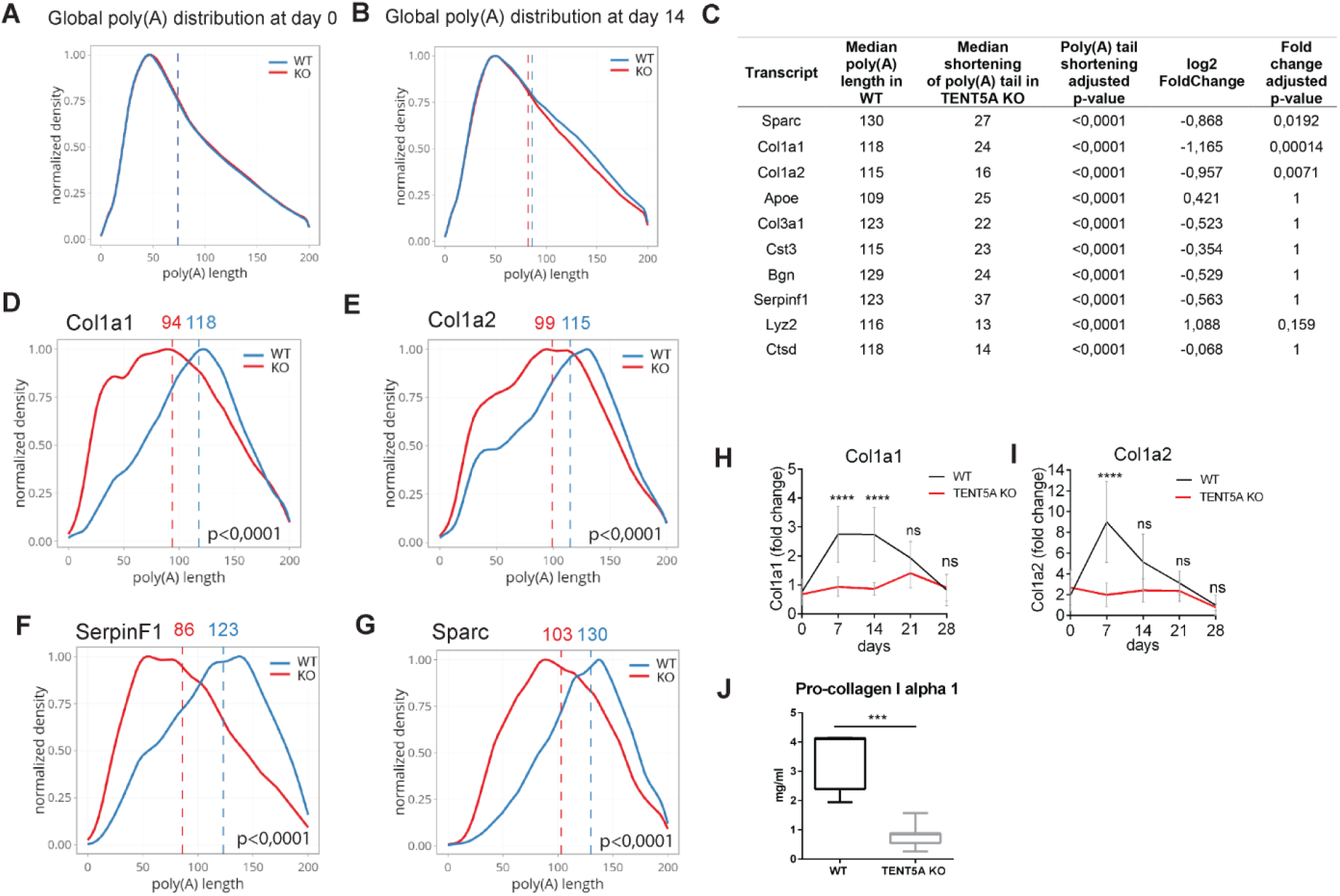
TENT5A polyadenylates and increases the expression of collagen I and other OI causative genes. A. DRS-based poly(A) length global profiling of mRNA isolated from WT and TENT5A KO neonatal calvarial osteoblasts at day 0 of maturation reveals no changes in poly(A) tail length. B. DRS-based poly(A) lengths global profiling of mRNA isolated from WT and TENT5A KO neonatal calvarial osteoblasts on day 14 of the maturation assay showing shortening of poly(A) tails in TENT5A KO mice. C. List of 10 transcripts with most shortened poly(A) tails in TENT5A KO D14 osteoblasts. Differential expression statistics were calculated with DESeq2 package. D-G: DRS-based poly(A) lengths profiling of Col1a1, Col1a2, SerpinF1 and Sparc mRNAs isolated from WT and TENT5A KO calvarial neonatal osteoblasts on day 14 of the maturation assay showing shortening of poly(A) tails in TENT5A KO osteoblasts. D. Col1a1: median poly(A) tail lengths (WT=118 nucleotides; TENT5A KO=94 nucleotides; p-value <0.0001). E. Col1a2: median poly(A) tail lengths (WT=115 nucleotides; TENT5A KO=99 nucleotides; p-value <0.0001). F. SerpinF1 (PEDF): median poly(A) tail lengths (WT=123 nucleotides; TENT5A KO=86 nucleotides; p-value <0.0001). G. Sparc (Osteonectin): median poly(A) tail lengths (WT=130 nucleotides; TENT5A KO=103 nucleotides; p-value <0.0001). H. RT-qPCR analysis of mRNA levels of Col1a1 during the osteoblast maturation assay, normalized to HMBS (n=5–6, medium values from technical triplicates) (p-values: D0, ns; D7, < 0.0001; D14, < 0.0001; D21, ns; D28, ns; Two-way ANOVA, Sidak’s multiple comparisons tests). I. RT-qPCR analysis of mRNA levels of Col1a2 during the osteoblast maturation assay, normalized to HMBS (n=4–6, medium values from technical triplicates) (p-values: D0, ns; D7, < 0,0001; D14, ns; D21, ns; D28, ns; Two-way ANOVA, Sidak’s multiple comparisons tests). J. Elisa measurement of the pro-collagen I alpha 1 level in WT and TENT5A serum (n=7; p-value = 0.0006, Mann–Whitney test). Data information: DRS (A-G) was performed in three biological replicates. Vertical dashed lines represent median poly(A) lengths for each condition.

Importantly, the poly(A) tails of the mRNAs of *Colla1* and *Colla2*, the most commonly mutated OI--causative genes, were noticeably shorter in TENT5A KO osteoblasts than in the WT. Median poly(A) tail length of *Colla1* mRNAs was decreased from 118 nucleotides in the WT to 94 nucleotides in the TENT5A KO (p < 0.0001, Fig 4D), whereas that of *Colla2* mRNA was decreased from 115 in the WT to 99 nucleotides in TENT5A KO (p < 0.0001, Fig 4E). Next, we examined the level of collagen I mRNAs through osteoblast maturation assay. In agreement with the results of DRS, both *Colla1* and *Colla2* mRNA levels were strongly decreased in TENT5A KO osteoblasts, especially on days 7 and 14 of mineralization (Fig 4H-I). Collagen deficiency was global and observed at the organismal level, as evidenced by the 75% decreased level of pro-collagen I alpha 1 in the serum of TENT5A KO mice compared with their WT littermates (Fig 4J).

The low level of collagen I in TENT5A KO mice is a plausible cause of the frequent bone fractures of OI type XVIII patients and TENT5A KO mice. However, their symptoms were more severe than those of OI type I patients and the Mov13 (+/-) mouse (Jaenisch et al., 1983), particularly with respect to bowing of the lower limbs, malformations, and short posture. This suggests that more factors may be involved in the pathogenesis of TENT5A--related OI. Interestingly in this respect, two other OI-causative genes: SPARC and SerpinF1 were also identified as TENT5A substrates by DRS and had the most shortened poly(A) tails among all TENT5A substrates. Both were downregulated at the protein level in TENT5A KO mice, presumably leading to a more severe phenotype (Fig EV3E-F).

Based on these results, we conclude that TENT5A polyadenylates Col1a1 and Col1a2 mRNA to increase its translation efficiency. This is the first study to report regulation of collagen I production at the post-transcriptional level by cytoplasmic modification of the length of its poly(A) tail. Moreover, we identified two additional substrates of TENT5A among OI-related genes, suggesting that the pathogenesis of type XVIII OI is complex.

### 5. Collagen I defect in TENT5A KO is both quantitative and qualitative

To determine whether the structure of collagen I fibers was compromised in TENT5A KO mice, we examined the migration patterns of collagen I extracted from the tendons of TENT5A WT and KO mice on SDS-PAGE gels. We did not detect any significant changes (Fig 5A), suggesting that posttranslational modifications of collagen I are not affected by TENT5A KO. This was confirmed by mass spectrometry analysis of SDS-PAGE collagen bands, which showed no differences in the global proline hydroxylation level or oxidation level (Fig 5B).

**Figure 5.**
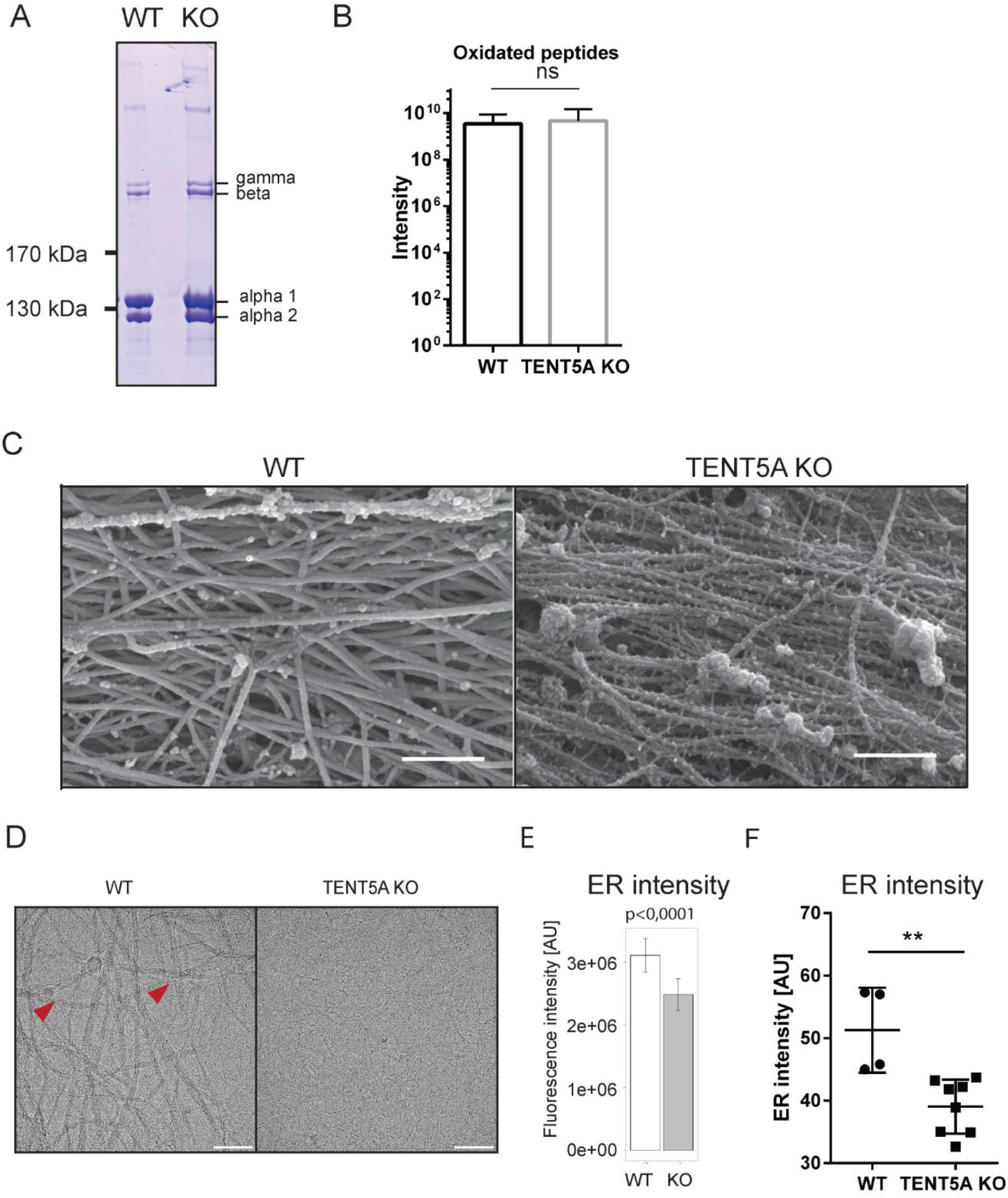
Lack of TENT5A leads to defects in collagen production A. Analysis of SDS-PAGE migration of collagen I isolated from WT (left) and TENT5A KO (right) tendons showing no differences between WT and TENT5A KO. B. Measurement of global proline hydroxylation using MS/MS. C. SEM images of collagen fibrils from mice femur shows prototypic fibrils in the TENT5A KO sample and compact collagen fibers in the WT sample. Scale bar, 500 nm. D. CryoEM visualization of isolated collagen I fibers from WT (left) and TENT5A KO (right) tendons. Arrowheads indicate fibers with a diameter of approximately 17.0±1.2 nm; fibers with a diameter of approximately 1.7±0.2 nm are observed as background. E–F. Endoplasmic reticulum is smaller in TENT5A than in WT adult long-bone derived osteoblasts. E. Osteoblasts were stained with anti-calreticulin antibody and fluorescence intensity was determined for 500 random cells. (p < 0.001; Unpaired t-test with Welsch correction). F. Osteoblasts were stained with ER tracker and analyzed by cell cytometry.

To examine collagen fiber structure *in situ*, we first performed scanning electron microscopy analysis of femur structure, which revealed a high level of disorder in collagen organization and assembly in TENT5A KO mice (Fig 5C). TENT5A KO fibrils were narrower than those of the WT (28.5 nm ±1.3 vs. 37.2 nm ±1.2, p < 0.001). Moreover, TENT5A KO fibrils exhibited disassembly into thinner prototypic fibrils of 11 nm ± 2nm, which were not observed in the WT. This characteristic differentiates collagen fiber robustness in the WT from those TENT5A KO, causing an abnormal and disarranged collagen fibers meshwork.

We next isolated collagen from mouse tendons using the acetic acid-pepsin method and visualized collagen by cryo-EM. This showed that WT tendon collagen preparation consisted of a mixture of two types of populations with diameters of 17.0±1.2 nm (red arrowhead) and 1.7±0.2 nm (observed as a background) whereas TENT5A KO tendon collagen consisted of one type with a diameter of 1.6±0.1 nm (Fig 5D). The finer diameter population corresponds to that of tropocollagen, while the thicker to that of fibril. The lack of fibrils in TENT5A KO preparation suggests that fibrils in these mice are extremely fragile and vulnerable to pepsin digestion.

As collagen I is the main protein secreted by osteoblasts, and the endoplasmic reticulum (ER) plays a crucial role in the process of osteogenesis, we measured the size of this subcellular organelle in TENT5A WT and KO osteoblasts. First, long bone-derived osteoblasts grown on glass coverslips were incubated with anti-calreticulin antibody to measure the area of the ER with respect to the whole-cell area delineated by HCS CellMask staining (Fig 5E). We also measured ER size by subjecting ER-tracker stained cells to flow cytometry (Fig 5F). Both approaches showed that the endoplasmic reticulum was smaller in TENT5A KO osteoblasts than in the WT.

Taken together, our results suggest that TENT5A KO animals have both quantitative and qualitative defects in collagen I production caused by aberrant mRNA polyadenylation of Col1a1, Col1a2, and collagen processing protein mRNAs.

### 6. Cytoplasmic adenylation plays a crucial role during osteoblast differentiation

We observed significantly longer poly(A) tails in samples derived from mineralizing osteoblasts (D14) than in those derived from non-differentiated neonatal calvarial osteoblasts (D0) from both WT and TENT5A KO osteoblasts (Fig 6A-B); however, the difference was clearer in the WT (median length of poly(A) tails: WT: D0, 74 nt; D14, 86 nt) than in the TENT5A KO (D0: 74 nt, D14: 82 nt). Since poly(A) tails were elongated not only in the WT but also in the TENT5A KO mice, upregulation of TENT5A expression during osteoblast differentiation may only be partially responsible. One possibility is that the expression of poly(A) polymerases other than TENT5A is upregulated during osteoblast differentiation. We found that, in addition to TENT5A, only TENT5C was upregulated (Fig 6C; EV4A). TENT5C KO mice do not exhibit any obvious skeletal dysplasia (Bilska et al., 2020; Mroczek et al., 2017; Zheng et al., 2019), but TENT5A/TENT5C KO mice showed preweaning lethality with almost complete penetrance (Fig 6D), suggesting redundancy in the poly(A) activities of TENT5A and TENT5C, possibly during bone formation.

**Figure 6.**
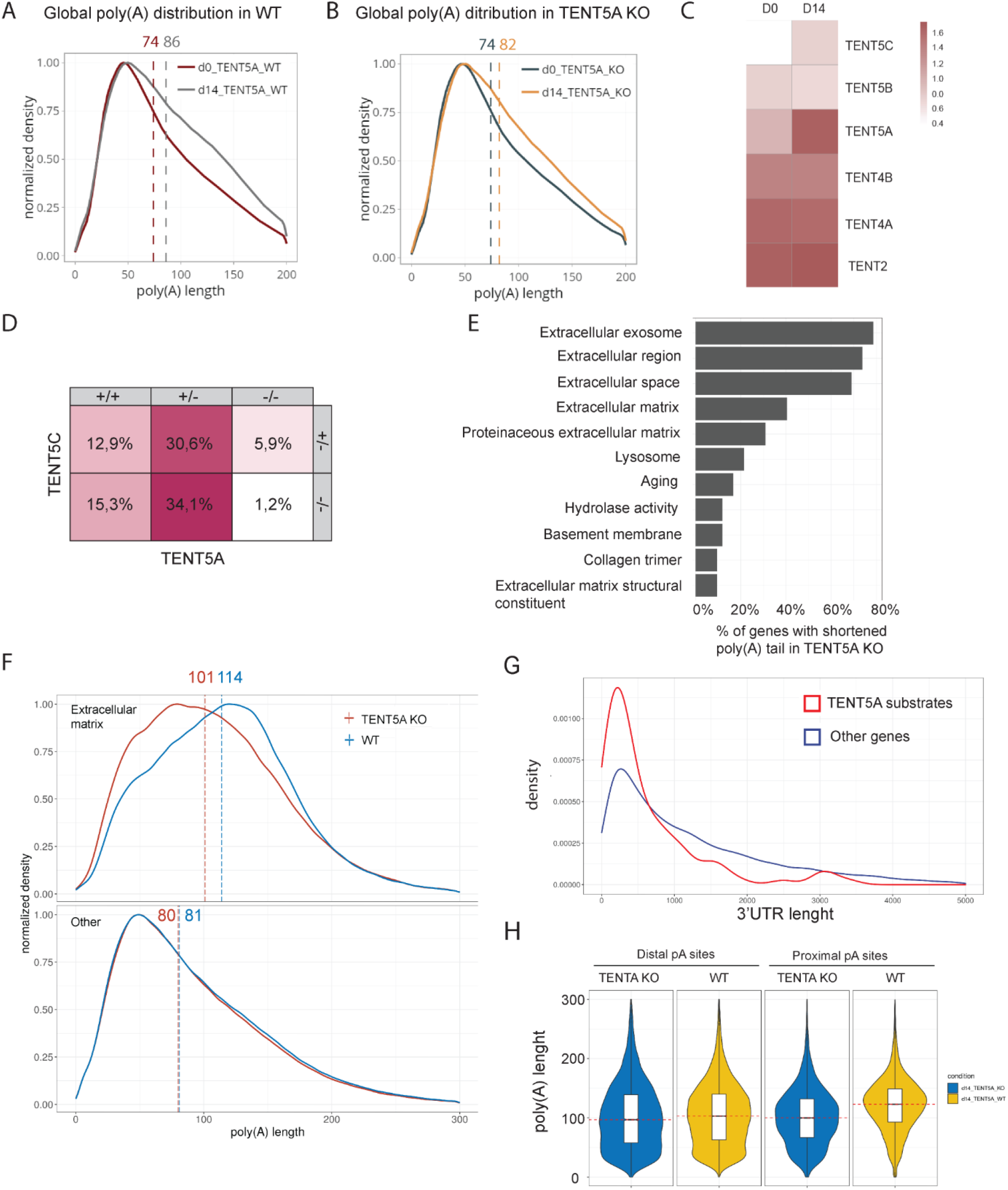
TENT5A is responsible for the wave of cytoplasmic polyadenylation that increases the expression of secreted proteins. A–B. DRC-based comparison of global poly(A) distribution in WT (A) and TENT5A KO (B) on day 0 and day 14 of the osteoblast maturation assay showing the existence of a polyadenylation wave during osteoblast differentiation, which was partially dependent on the activity of TENT5A. A. WT: median poly(A) tail lengths at D0 = 74 nucleotides, at D14 = 86 nucleotides B. TENT5A KO: median poly(A) tail lengths at D0 = 74 nucleotides, at D14 = 82 nucleotides C. Heatmap showing expression of non-canonical poly(A) polymerases in D0 and D14 osteoblasts. The only poly(A) polymerases upregulated during osteoblast differentiation were TENT5A and TENT5C. D. Preweaning lethality of TENT5A/C KO with incomplete penetrance. Observed frequency of TENT5A(-/-); TENT5C (-/) is 1.2% instead of the expected 12.5% and TENT5A(-/-); TENT5C(WT/-) is 5.9% instead of the expected 12.5%. n=85; p = 0,0065; Chi-square test E. Functional GO term annotation of transcripts with shortened poly(A) tails in TENT5A KO. F. Distribution of poly(A) tail lengths of mRNA encoding extracellular matrix proteins (top) and other proteins (bottom) in WT and TENT5A KO neonatal calvarial osteoblasts on D14 of the maturation assay. Extracellular matrix mRNA medium poly(A) tail length for WT =114 nucleotides; for TENT5A KO =101 nucleotides; other mRNAs: WT = 81 nucleotides; TENT5A KO = 80 nucleotides. G. Metagene analysis showing the distribution of 3’UTR lengths in TENT5A substrates and other mRNAs of neonatal calvarial osteoblasts at D14 of the maturation assay. H. Violin plot showing the distribution of poly(A) tail lengths of different Col1a2 isoforms arising from alternative polyadenylation sites.

To determine the specificity of TENT5A, we performed gene ontology analysis, which revealed that transcripts encoding secreted proteins and especially extracellular matrix constituents were strongly enriched among transcripts with shortened poly(A) tails in TENT5A KO mice (Fig 6E). These transcripts have relatively long poly(A) tails and seem to be predominant substrates of TENT5A since their exclusion from the analysis eliminates the global difference in poly(A) tails lengths between WT and TENT5A KO on D14 (EV4B Fig 6F,). To see if the effect also applies to osteoblasts isolated from long-bones of adult animals, we performed an additional DRS experiment. Although the results were less reproducible, the distribution of poly(A) tails lengths was strikingly similar to that of mRNAs in mineralizing (D14) osteoblasts isolated from neonatal calvaria. In other words, the average lengths of poly(A) tails were relatively long, and mRNAs encoding extracellular matrix constituents, including Col1a1 and Col1a2, were TENT5A substrates (Fig EV5A-D). Western blot analysis of cellular osteoblastic fractions revealed that membranes are enriched in TENT5A protein, which is in agreement with the localization of its substrates to the ER (Fig EV4C)

To determine what distinguishes TENT5A substrates, we first performed 3’UTR motif analysis of transcripts identified as TENT5A substrates. No specific motive was found, despite the canonical polyadenylation signal (Fig EV4D) Nonetheless, TENT5A substrate mRNA was relatively short, had a higher %GC content, and tended to have short 3’UTRs (Fig 6G, EV4E-F). We then searched for potential alternative polyadenylation sites among TENT5A-responsive genes, which revealed that several collagens are alternatively polyadenylated. Importantly, in all cases, mRNAs with shorter 3’UTRs employing a proximal poly(A) signal were more TENT5A dependent and had longer poly(A) tails. In Col1a1 and Col1a2 transcripts, which possess two alternative polyadenylation sites, the proximal pA site was clearly more responsive to TENT5A (Fig 6H).

In conclusion, our analysis reveals that TENT5A, possibly in cooperation with TENT5C, is responsible for the cytoplasmic polyadenylation of extracellular matrix constituents. It also for the first time suggests the existence of a wave of cytoplasmic polyadenylation occurring during osteoblast mineralization mainly focused on ER-targeted mRNAs.

## Discussion

In this paper, we describe a post-transcriptional mechanism essential for proper bone formation. TENT5A polyadenylates and enhances the expression of proteins secreted by osteoblasts that are crucial for the mineralization process. Patients suffering from type XVIII OI display severe symptoms of the disease, which include numerous spontaneous fractures appearing as soon as early infancy, congenital bowing of lower limbs, hypomineralization, and in some cases death in childhood (Doyard et al., 2018). Type XVIII patients also have blue sclerae, which is consistent with the observed in TENT5A KO mice defect in collagen I production. The severe phenotype of type XVIII patients can be explained by the relatively broad spectrum of TENT5A substrates identified in the TENT5A KO mouse model, which recapitulates the disease symptoms of type XVIII OI. TENT5A targets mRNAs not only encoding collagen chains and other proteins directly involved in collaged secretion/folding (EV Table 1, EV Table 2), but also SerpinF1 mRNA, whose expression was clearly downregulated in TENT5A KO mice. Mutation of SerpinF1, which encodes a secreted collagen-binding protein that participates in various signaling pathways, is responsible for type VI OI, whose characteristic feature, elevated ALP levels, was also observed in type XVIII OI. However, because of the diversity of TENT5A regulated mRNAs, it is impossible at present to describe in detail how downregulation of a particular TENT5A substrate contributes to diseases syndromes.

We provide evidence that during osteoblast differentiation, TENT5A and TENT5C are induced and responsible for a wave of polyadenylation of secreted proteins. This is analogous to the role of TENT5C in B cells, whose main substrates are immunoglobulins. In osteoblasts, as indicated above, the range of TENT5A substrates is more divergent than those of TENT5C in B cells. The mechanism that determines substrate specificity has yet to be identified; however, it is clear that TENT5A and TENT5C perform a novel type of cytoplasmic polyadenylation that is different from the waves of polyadenylation previously described during oocyte maturation and postulated to be involved in neuronal processes (Villalba et al., 2011; Wu et al., 1998). TENT5A/TENT5C substrates do not undergo translational inactivation via deadenylation and are not stored, but are probably actively engaged in translation at the endoplasmic reticulum, where TENT5A/TENT5C poly(A) polymerases are enriched. Moreover, no specific sequence motive or other sequence feature, apart from relatively short UTRs and a preference for proximal poly(A) sites, were found in TENT5 substrates, which is in contrast to CPEB-mediated regulation of polyadenylation in oocytes (Ivshina et al., 2014)

Finally, our results change the generally held view of poly(A) tail metabolism. In osteoblasts, the most abundant transcripts have long poly(A) tails, which is in sharp contrast to recent data indicating that the lengths of poly(A) tails of highly abundant mRNAs are relatively short (Lima et al., 2017). Clearly, the dynamics of poly(A) tail metabolism differ according to cell type and organ, and results obtained using one particular model system cannot be extrapolated to other systems. Our knowledge of poly(A) tail metabolism is still very fragmentary and there is much to be discovered in the future.

## Methods

### Mice

TENT5A knock-out mouse lines with a loss of function deletion in exon 2 (c.436_462del26) in C57BL/6N genetic background and (c.403_406del4ins50bp) in C57BL/6/Tar x CBA/Tar mixed background, were established using the CRISPR/Cas9 method. TENT5A-3xFLAG knock-in was established in C57BL/6/Tar x CBA/Tar mixed background mice. TENT5C mouse line was established previously (Bilska et al., 2020; Mroczek et al., 2017). Experimental mice originated from heterozygotic matings and were cohoused with littermates. TENT5A KO (C57BL/6/Tar x CBA/Tar) and TENT5A-3xFLAG mice were bred in the animal house of Faculty of Biology, University of Warsaw. TENT5A KO (C57BL/6/Tar x CBA/Tar), TENT5A-3xFLAG, TENT5C KO, and TENT5C-3xFLAG mice were maintained under conventional conditions in open polypropylene cages filled with wood chip bedding (Rettenmaier). The environment was enriched with nest material and paper tubes. Mice were fed *ad libitum* with a standard laboratory diet (Labofeed B, Morawski). In rooms, humidity was kept at 55± 10% and temperature at 22°C ±2 °C, with 12 h/12 h light/dark cycles (lights were on from 6:00 to 18:00) and at least 15 air changes per hour. TENT5A KO (C57BL/6N) mice were bred at Czech Center of Phenogenomics and maintained in individually ventilated cages in a room with controlled temperature (22±2 °C) and humidity under a 12 h light/12 h dark cycle. Food (Standard diet from Altromin) and drink were provided *ad libitum*. Animals were closely followed-up by the animal caretakers and researchers, with regular inspection by a veterinarian, according to the standard health and animal welfare procedures of the local animal facility. No statistical method was used to predetermine sample size. All animal experiments were approved by the Animal Ethics Committee of the Czech Academy of Sciences (primary screen project number: 62/2016 and secondary screen project number: 45/2017) or by the Local Ethical Committee in Warsaw affiliated to the University of Warsaw, Faculty of Biology (approval numbers: 176/2016, 732/2018, decision number 781/2018) and were performed according to Czech guidelines for the Care and Use of Animals in Research and Teaching or according to Polish Law (Act number 266/15.01.2015).

### Mouse Genotyping

DNA isolation was performed using the HotShot method (Truett et al 2000) with minor modifications. For ear and tail tips from mature and newborn mice, the volumes of alkaline and neutralization solutions were scaled up to 175 μl and 100 μl, respectively. Lysis time was reduced to 30 minutes. Crude DNA extract (1 μl) was added to 19 μl of PCR mix containing Phusion HSII polymerase, HF buffer (Thermo), and 10pM of primers (Tent5A_seq1F and Tent5A_seq1R for genotyping of TENT5A KO mice and Tent5A_seq1F and Tent5A_seq1R for genotyping TENT5A-3xFLAG mice; Appendix Table 1). Genotyping of TENT5C KO mice was performed as described previously (Bilska et al., 2020).

### Strains used

The following experiments were performed using TENT5A KO mice with the C57BL/6J/Tar × CBA/Tar mixed background: initial phenotyping, DRS, osteoblast maturation assay, consequential analysis whole mount staining, collagen migration analysis, cryo-EM, pro-collagen I level analysis in serum, western blot analysis, mass spectrometry analysis, μCT, osteoblast proliferation analysis, and SEM. The following experiments were performed using TENT5A KO mice with the C57BL/6N background: initial phenotyping, μCT, analysis of serum biochemistry, and SEM.

### Analysis of lethality

For TENT5A KO mice, analysis were performed independently on day 6 and day 35 on mice derived from TENT5A(WT/-) × TENT5A(WT/-) matings. Mice were genotyped as described above.

For TENT5A/TENT5C dKO analysis was performed retrospectively on 85 weanling mice deriving from TENT5A(WT/-); TENT5C(-/-) × TENT5A(WT/-); TENT5C(WT/-) matings. This choice of mating strategy was made based on animal welfare and fertility issues.

### Whole-Mount Skeletal Staining

Alizarin Red/Alcian Blue staining of E17, P6, and adult WT and TENT5A KO mouse was performed as described previously (Hilton, 2014).

### Immunohistochemistry

For immunodetection of FLAG in bones, femurs from 11-week-old mice were fresh-frozen in Killik medium (Bio-Optica, Milan, Italy) and 10-μm sections were cut using the Kawamoto method (Kawamoto, 2003) on adhesive films (Section-Lab, Hiroshima, Japan) with a cryostat (Leica CM 1950, Leica, USA).

Sections on films were fixed in 4% PFA for 10 minutes at 4 °C, washed in TBS, incubated with Proteinase K for antigen unmasking (5 μg/ml in TBS, 10 minutes at room temperature (RT)), quenched with 3% H2O2 in TBS, and washed again with TBS. Then, sections were blocked with 10% donkey serum, 1% BSA and 0.3% Triton X-100 in TBS and incubated overnight with rabbit anti-FLAG antibody (Appendix Table 2). The next day, sections were washed with wash buffer (TBS with Triton X-100 0.025%), incubated with donkey anti-rabbit IgG antibody conjugated with HRP (Agrisera, Vannas, Sweden; diluted 1:200 in blocking solution) for 1 h at RT together with Hoechst 33342 diluted 1:1000. After incubation, sections were washed and developed in CF488A-tyramide (Biotium, Fremont, CA, USA) diluted 1:100 in 0.1M borate buffer pH 8.7 with 0.1% Tween-20 and 0.003% H2O2 (5’ at RT), washed with TBS, and mounted on microscope slides with Prolong Gold (Invitrogen).

Stained specimens were scanned using an Opera Phenix high-throughput confocal system (PerkinElmer, Waltham, MA, USA) equipped with a 40 × water immersion objective. The obtained tile arrays of stack series were used for flatfield correction with BaSiC Tool (Peng et al., 2017) at the default settings, maximum orthogonal projection and stitching with Grid/Collection Stitching Plugin (Preibisch et al., 2009), all using (Fiji Is Just) ImageJ software.

### Serum biochemical analysis

Biochemical data were collected from the phenotyping pipeline performed by the International Mouse Phenotyping Consortium (IMPC) in the Czech Center for Phenogenomics (CCP). Sixteen-week-old mice were used in experiments, and nine TENT5A KO males and six TENT5A KO females were analyzed against a WT cohort. Blood samples were taken from isoflurane-anesthetized mice by retro-bulbar sinus puncture with non-heparinized glass capillaries. Samples were collected in lithium/heparin-coated tubes (KABE cat # 078028). After collection, each sample was mixed by gentle inversion and then kept on RT until centrifugation. Samples were centrifuged within 1 h of collection at 5000 × g, for 10 minutes at 8 °C. Once separated from the cells, plasma samples were analyzed using a Beckman AU480 biochemical analyzer.

### Mouse body weight analysis

Five-week-old littermates deriving from heterozygotic matings were weighed using a standard laboratory scale.

### Preparation of long bones

Tibia and fibula were dissected and boiled in 100 °C water for 5 h without stirring. Soft tissues were separated from the bone. Pre-cleaned bones were subjected to 30% hydrogen peroxide treatment and incubated for 24–48 h. The obtained preparations were washed with PBS, photographed, and stored dry.

### MicroCT scanning and analysis

Five mice from each genotype (WT and TENT5A KO) were sacrificed by cervical dislocation at 13–14 weeks of age. Mice in 4% PFA were transported to the Czech Center for Phenogenomics (CCP) for μCT analysis of the whole skeleton and high-resolution analysis of femurs.

First, the whole body scan was performed in a SkyScan 1176 instrument (Bruger, Belgium) at a resolution of 9 μm per voxel (0.5 mm Al filter; voltage, 50 kV; current, 250 μA; exposure, 2000 ms; rotation, 0.3°; spiral scan, 2x averaging) in a wet atmosphere. Reconstruction was performed in an NRecon 1.7.1.0 (Bruker, Belgium) with the following parameters: smoothing = 2, ring artifact correction = 3, beam hardening correction = 36%, and defect pixel masking threshold = 10%. The range of intensities was set from 0.004 AU to 0.23 AU.

Then, femur bones were extracted from the mice and mounted in 2.5% low melting agarose (Sigma-Aldrich Co., USA). After at least 1 day in the fridge (4 °C) for sample stabilization, they were scanned in a SkyScan 1272 (Bruker, Belgium) at a resolution of 1.5 μm per voxel (Al filter, 1 mm; voltage, 80 kV; current, 125 μA; exposure, 2584 ms, rotation, 0.21° in a 360° scan, 2x averaging). NRecon 1.7.3.1 (Bruker, Belgium) with the InstaRecon 2.0.4.0 (InstaRecon, USA) reconstruction engine was used to obtain digital sections. Reconstruction was performed with the following parameters: smoothing = 6, ring artifact reduction = 8, beam hardening correction = 28%, and defect pixel masking threshold = 10%. The range of intensities was set from 0.00 AU to 0.110 AU.

Reconstructions were reoriented to the same orientation in DataViewer 1.5.4.0 (Bruker, Belgium) and subsequently, regions of interest for trabecular and cortical bones analysis were selected in CT analyzer 1.18.4.0 (Bruker, Belgium). Bone was separated from the background by the Otsu method (CIT). Regions of interest for trabecular bone were selected automatically based on the Bruker Method note MCT-124 with some modifications due to the high resolution of the scan. Parameters, such as bone volume, porosity, and bone mineral density (BMD) in cortical bone, and relative bone volume, volume:surface ratio, structure linear density, orientation, and thickness in trabecular bone, were measured. BMD was established with calibrated Hydroxyapatite (HAP) phantoms (25% and 75%) scanned and reconstructed under the same conditions as for samples. CTvox 3.3.0 (Bruker, Belgium) was used for scan visualization and image processing.

### Neonatal murine calvarial osteoblast isolation

Primary osteoblast cultures were established using the standard collagenase method (Hilton, 2014). Briefly, three to six old neonates from heterozygotic matings were euthanized using isoflurane and decapitated. Mice were genotyped, and calvaria were isolated and pooled (four per culture), rinsed with PBS, and subjected to five rounds of digestion using type II collagenase (Thermo Fisher Scientific, 17101015). Digests three to five were collected for culture. All osteoblast cultures were grown in MEM alpha medium supplemented with 10% FBS (Sigma) and penicillin-streptomycin (Thermo Fisher Scientific).

### Osteoblast maturation assay

Osteoblasts were isolated as described above and cultured until they reached confluency. Cells were seeded into 12-wells plates at 35000 cells/well. After reaching confluency, medium was changed to MEM alpha supplemented with 10% FBS, 50 μg/ml sodium ascorbate (Sigma), and 10 mM β-glycerophosphate (Roth).

Cells were collected and stained on days 0, 7, 14, 21, 28, and 35. Medium was changed every 2–3 days. Cell were detached using trypsin (ThermoFisher Scientific). For RNA isolation collected cells were washed with PBS and resuspended in TRI reagent (Sigma). For protein extraction cells were resuspended in PBS containing 0.1% NP-40 and protease inhibitors and incubated for 30 minutes in 37 °C in the presence of 250 U of Viscolase (A&A Biotechnology).

For staining cells were fixed in 4% formaldehyde and washed either three times with distilled water (for NBT/BCIP) or three times with PBS and once in 96% ethanol (for Alizarin Red). Staining was performed using 0.1% Alizarin Red S (Sigma) in 95% ethanol or NBT/BCIP Substrate Solution (ThermoFisher Scientific).

### Western blot analysis

For western blot analysis, equal amount of cells were lysed in PBS containing with 0.1% NP40, protease inhibitors, and viscolase (A&A Biotechnology, 1010-100) for 30 minutes at 37°C. After shaking at 600 rpm and homogenization with a Dounce homogenizer, Laemmli buffer was added and samples were denatured for 10 minutes at 100 °C. Samples were separated on 10–12% SDS-PAGE gels and proteins were transferred to Protran nitrocellulose membranes (GE Healthcare), after which membranes were stained with 0.3% w/v Ponceau S in 3% v/v acetic acid and digitized. Membranes were incubated with 5% milk in TBST buffer for 1 h followed by overnight incubation in 4 °C with specific primary antibodies (Appendix Table 2).

### Immunostaining of osteoblasts

Neonatal osteoblasts were isolated as described above and at P1 were seeded onto glass 18-mm coverslips in a 12-well plate. The next day, cells were fixed using paraformaldehyde (10 minutes in 4% in 0.1M phosphate buffer, pH 7.4) and washed with PBS. For immunodetection of FLAG, fixed cells on coverslips were treated in the same way as tissue cryosections on film, as described the Immunohistochemistry section, but without the antigen unmasking step. Stained cells were imaged using a LSM800 confocal microscope (Zeiss, Jena, Germany) equipped with 20× air and 63 × oil immersion objectives. Collected Z-stacks were used to generate orthogonal maximum projections using ImageJ software.

### RNA isolation

Total RNA was isolated using TRIzol (Thermo Fisher Scientific) according to the manufacturer’s instructions, dissolved in nuclease-free water, and stored at −80 °C.

### RT-qPCR

For quantitative analysis, RNA was first treated with DNase (TURBO DNA-free Kit, Invitrogen; AM1907) for 30 minutes at 37 °C and then reverse transcribed using SuperScript III (Invitrogen; 18080085), oligo(dT)20, and random-primers (Thermo Fisher Scientific). Quantitative PCR was performed using Platinum SYBR Green qPCR SuperMix-UDG (Thermo Fisher Scientific; 11733046) in a LightCycler 480 II (Roche) PCR device and the primers listed in Appendix Table 1. Gene expression was normalized to that of HMBS (Stephens et al., 2011). Differences were determined using the 2^-ΔΔC(t)^ calculation.

### Osteoblast isolation from murine adult long bones

Isolation of osteoblasts from adult long bones was performed as described previously with minor modifications (Bakker and Klein-Nulend, 2012). Briefly, febur and tibia were isolated, and muscles and surrounding tissue were removed. Bone marrow was removed by centrifugation. Diaphyses were cut into small pieces and bone pieces were washed several times with PBS solution. Bone pieces were incubated in OptiMEM medium (Thermo Fisher Scientific) containing 1 mg/ml collagenase II (Thermo Fisher Scientific) for 2 h at 37 °C in a shaking water bath. Bone pieces were rinsed several times with DMEM (Thermo Fisher Scientific) containing 10% FBS (Sigma), and then transferred to T25 flask containing the same medium. Cell were cultured for 10–14 days and medium was changed two or three times per week. Experiments were performed at passage 3.

### Cell proliferation analysis

Adult osteoblasts were isolated as described above. To calculate the proliferation ratio, cells were stained with 1 μM CFSE according to the manufacturer’s instructions. The signal was measured at time 0 h, 48 h and 96 h, and qMFI were calculated. The proliferation ratio was calculated based on the gMFI. The intensity of fluorescence was measured as gMFI. Cells were measured with BD LSRFortessa™ under FACS Diva Software v8.0.1 (BD) software control and analyzed using FlowJo (Data Analysis Software v10).

### Nanopore direct RNA sequencing (DRS)

#### Cell culture and RNA retrieval

Neonatal calvarial and adult long-bones osteoblasts were isolated as described above. Adult osteoblasts were passaged three times before harvesting. Neonatal osteoblasts were passage once. After reaching confluency, cells were either harvested (for D0 timepoint) or medium was changed to differentiation medium (MEM alpha supplemented with 10% FBS, 50 μg/ml sodium ascorbate (Sigma) and 10 mM β-glycerophosphate (Roth) and the cells were cultured for 14 days with two medium changes per week and harvested (for the D14 time point). RNA was isolated as described before. The cap-enriched mRNA was prepared from 100 μg of total RNA with GST-eIF4E^K119A^ protein and glutathione sepharose 4B (GE Healthcare), as described previously (Bilska et al., 2020).

#### Library preparation and sequencing

DRS libraries were prepared using Direct RNA Sequencing (ONT, SQK-RNA002) with 5 μg of murine cap-enriched mRNA according to the manufacturer’s instructions, and to optimize sequencing efficiency, were mixed with 100–150 ng of *Saccharomyces cerevisiae* oligo(dT)-enriched mRNA. Sequencing was performed using a MinION device, MinKNOW 19.10.1 software, Flow Cell (Type R9.4.1 RevD), and basecalling with Guppy 3.3.0 (ONT). Raw sequencing data (fast5 files), as well as basecalled reads were deposited at ENA (project accession number: PRJEB39819). Summary of sequencing runs is presented in the Appendix Table 5.

#### Bioinformatic analysis

Obtained reads were mapped to GencodeVM22 reference transcript sequences (Frankish et al., 2019) using Minimap 2.17 (Li, 2018), with options -k 14 -ax map-ont-secondary=no and processed with samtools 1.9 to filter out supplementary alignments and reads mapping to the reverse strand (samtools view -b -F 2320). The poly(A) tail lengths for each read were estimated using Nanopolish 0.13.2 polya function (Li, 2018). In subsequent analyses, only length estimates with QC tag reported by Nanopolish as PASS were considered. Statistical analysis was performed using functions provided in the NanoTail R package (https://github.com/smaegol/nanotail, manuscript in preparation). In detail, the Generalized Linear Model approach, with log2(polya length) as a response variable, was employed, and transcripts that had a low number of supporting reads in each condition (<20) were filtered out. To correct for the batch effect, a replicate identifier was used as one of the predictors, in addition to the condition (Tent5A KO/WT) identifier. P values (for the condition effect) were estimated using the Tukey HSD post hoc test and adjusted for multiple comparisons using the Benjamini–Hochberg method. Transcripts were considered as having a significant change in poly(A) tail length, if the adjusted P value was < 0.05, the absolute value of calculated Cohen’s d (effect size) was >0.2, and there were at least 20 supporting reads for each condition. Functional Enrichment Analysis was done using DAVID Functional Annotation Tool (Huang et al., 2009a, 2009b)

For differential expression estimates, reads were mapped to the mouse GRCm38 genome using Minimap 2.17 (Li, 2018), with options -k 14 -ax splice -uf, Features were assigned using Gencode VM22 and featureCounts from the subread package (Li, 2018) in the long read, strand-specific mode (-L -s 1), including only features covered by at least 20% (--fracOverlapFeature 0.2) and reads overlapping with a feature by at least 50% (--fracOverlap 0.5). Statistical analysis of differential expression was performed using the DESeq2 (v.1.24.0). Bioconductor package (Li, 2018), using default settings and correcting for the batch effect.

### Motif enrichment analysis

Fasta sequences of 3’UTRs of (1) TENT5A substrates and (2) all Gencode-annotated transcripts in mm10 genome (background) were obtained using bedtools getfasta tool (v. 2.29.2) (Quinlan and Hall, 2010), using bed files with 3’UTR coordinates downloaded from UCSC Table Browser tool (GENCODE VM23 track and known_gene table) (Karolchik et al., 2004) and GRCm38 genome sequence. Sequence motifs enriched in 3’UTRs of TENT5A substrates were identified using DREME tool (Bailey, 2011), run with options -rna -norc -k 8 -l, with background set to 3’UTRs of all Gencode-annotated transcripts in mm10 genome.

### Measurement of pro-collagen I level

Elisa for pro-collagen I was performed using Mouse Pro-Collagen I alpha 1 ELISA Kit (Abcam, ab210579) according to manufacturer’s instructions with 1:2000–1:4000 serum dilutions.

### Tethering assay

Tethering assays were performed as previously described (Chekulaeva et al., 2011). In brief, one day before transfection, 0.75 ml of HEK293 cells were seeded into 6-well plates to achieve about 70–80% confluence on the day of transfection. Next, cells were co-transfected with 100 ng of constructs expressing the reporter Renilla luciferase (RL-5BoxB), 100 ng of control firefly luciferase (FL, pGL3 plasmid), and 2 μg of plasmid encoding tethered NHA-protein using 5 μl of Lipofectamine 2000 and OPTI-MEM media (Invitrogen) according to manufacturer’s instructions. All transfections were repeated at least three times.

### Northern blot

Low-molecular weight RNA samples were separated on 4–6% acrylamide gels containing 7M urea in 0.5× TBE buffer and transferred to a Hybond N+ membrane by electrotransfer in 0.5× TBE buffer. High-molecular weight RNA samples were separated on 1.2% agarose gels in 1× NBC buffer containing formaldehyde and transferred to membranes by capillary elution using 8× SSC buffer crosslinked by 254 nm UV light. Radioactive probes were prepared with a DECAprime II DNA Labeling Kit (Invitrogen) according to manufacturer’s instructions. Northern blots were carried out in PerfectHyb Plus Hybridization Buffer (Sigma), scanned with Fuji Typhoon FLA 7000 (GE Healthcare Life Sciences), and processed with Multi Gauge software Ver. 2.0 (FUJI FILM).

### Collagen isolation from tendon

For Cryo-EM and migration analysis, collagen from tendons was isolated using the acetic acid/pepsin method as described previously (Pokidysheva et al., 2013). Briefly, tendons were isolated from 9–10-week-old WT and TENT5A mice. After an initial 4 h incubation in 0.5 M acetic acid, tissues were digested in 0.5 M acetic acid containing 1 mg/ml porcine pepsin (Sigma) for 20 h. Samples were centrifuged to remove insoluble material and NaCl was added at a final concentration of 0.7 M to precipitate collagens. After 2 h of incubation, samples were centrifuged (20000 × g, 1 h, 4 °C) and precipitates were resuspended in 0.1 M acetic acid. After pH neutralization using 1 M Hepes pH 8 (Sigma), NaCl was added at a final concentration of 2.5 M and collagen was precipitated for 7 h. After centrifugation (20000 × g, 1 h, 4 °C) precipitates were resuspended in 0.1 M acetic acid to achieve a collagen concentration of approximately 1 mg/ml.

### Collagen migration analysis

Protein concentration in isolated collagen samples (described above) was measured at 280 nm using Nanodrop OneC (Thermo Scientific). After pH neutralization using 1 M Hepes pH 8 (Sigma), samples were diluted in Laemmli sample buffer and 2 μg of protein was separated on a NuPage 3–8% Tris-Acetate gel (Invitrogen). The gel was then stained with Coomassie Blue and digitalized.

### Mass Spectrometry

Collagen was isolated and separated as described above. Bands corresponding to Col1a1 and Col1a2 were cut into slices and subjected to the standard “in-gel digestion” procedure, during which proteins were reduced with 100 mM DTT (for 30 minutes at 56 °C), alkylated with 0.5 M iodoacetamide (45 minutes in a darkroom at RT), and digested overnight with 10 ng/μl trypsin solution (sequencing grade modified trypsin, Promega V5111). The resultant peptides were eluted from the gel with 0.1% trifluoroacetic acid (TFA) and 2% acetonitrile (ACN). Finally, to stop digestion, trifluoroacetic acid was added at a final concentration of 0.1%. The digest was centrifuged at 14 000 × g for 30 minutes at 4 °C to pellet solids. The particle-free supernatant was analyzed by LC-MS/MS in the Laboratory of Mass Spectrometry (IBB PAS, Warsaw) using a nanoAcquity UPLC system (Waters) coupled to an Orbitrap QExative mass spectrometer (Thermo Fisher Scientific). The mass spectrometer was operated in the data-dependent MS2 mode, and data were acquired in the m/z range of 300–2000. Peptides were separated on a 180 min linear gradient of 95% solution A (0.1% formic acid in water) to 35% solution B (acetonitrile and 0.1% formic acid). Each sample measurement was preceded by three washing runs to avoid cross-contamination. The final MS washing run was searched for the presence of cross-contamination between samples.

Data were searched with Max-Quant (Version 1.6.3.4) platform search parameters: match between runs (match time window, 0.7 minutes; alignment time, 20 minutes); enzyme, trypsin/p specific; max missed, 2; minimal peptide length, 7 aa; variable modification, methionine and proline oxidation; fixed, cysteine alkylation; main search peptide tolerance, 4.5 ppm; protein FDR, 0.01. Data were searched against the protein database containing all mice collagen sequences.

The list of peptides containing oxidized proline was statistically analyzed. The list of peptides containing oxidized proline was statistically analyzed. No difference has been detected both at the level of a single peptide and at the general level of sum of a intensities of all oxidated peptides identified in WT and TENT5A KO samples.

### Femur collagen visualization in SEM

Seven-week-old mice femurs were dissected and transferred to 0.9% saline physiological solution. Epiphysises were cut off, and bones were flushed out. Bone tissue was cut into 2-mm pieces and placed in PBS buffer containing 4% paraformaldehyde (Schuchardt, Muenchen, Germany) and 1% glutaraldehyde (Merck, electron microscopy grade, Sigma-Aldrich, Czech Republic) for 1 h at RT and then 1 week at 4 °C. Fixed bone tissue pieces were extensively washed on a rotator (PBS buffer, three times, 20 minutes, RT) and post-fixed in 1% OsO4 (1 h, RT). Post-fixed samples were extensively rewashed on a rotator (ddH2O, three times, 20 minutes, RT) and dehydrated in the graded alcohol series (25%, 50%, 75%, 90%, 96%, 100% and 100%, 20 minutes each). Finally, the samples were critical point dried (K850 Critical Point Dryer, Quorum Technologies Ltd, Ringmer, UK).

Dried tissue pieces were mounted onto standard 12.5-mm aluminum stubs (Agar Scientific, UK) using Ultra Smooth Carbon Discs (SPI Supplies, USA) or with Silberleitlack (Ferro GmbH, Frankfurt am Main, Germany). The samples were then sputter-coated with 3 nm of platinum (Turbo-Pumped Sputter Coater Q150T, Quorum Technologies Ltd, Ringmer, UK).

The mounts were examined in a FEI Nova NanoSEM 450 field emission gun scanning electron microscope (FEI, Brno, Czech Republic, now Thermo Fisher Scientific) at 3–5 kV using ETD, CBS, and TLD detectors. Sample charging, when it occurred, was eliminated in the beam deceleration mode of the scanning electron microscope.

### Cryoelectron microscopy

Collagen preparations were plunge-frozen onto Quantifoil R2/2 holey carbon grids using a Thermo Fisher Vitrobot. CryoEM data collection was performed using a Thermo Fisher Glacios TEM operating at 200 kV, equipped with a 4k × 4k Falcon 3EC direct electron detection camera at a magnification of 92k, corresponding to a pixel size of 1.5 Å at the specimen level.

### Osteoblast fractionation

Neonatal osteoblasts were isolated as described above. Cells were harvested at passage 1, and fractionation was performed using Subcellular Protein Fractionation Kit (Thermo Fisher Scientific). In parallel, total protein fractions were prepared by lysing cells in PBS containing 0.1% NP40, protease inhibitors (Invitrogen), and Viscolase (A&A Biotechnology).

### ER size analysis

Adult long-bone derived osteoblasts were isolated as described above.

For microscopic analysis, osteoblasts were seeded onto 18-mm glass coverslips in a 12-well plate and fixed with 4% formaldehyde (Sigma) the next day. Fixed cells were permeabilized using ice-cold methanol in −20 °C for 10 minutes and blocked in blocking buffer (PBS with 5% goat serum and 0.3% Triton X-100) for 30 minutes. Incubation with primary anti-calreticulin antibody (Cell signaling #12238; 1:600; O/N; 4 °C, wet chamber) and secondary goat anti-rabbit HRP conjugated antibody (Agrisera, AS101069; 1 h, RT) was performed in antibody dilution buffer (1% BSA, 0.1% Triton X-100 in PBS). Cells were stained with Hoechst 33342 (Thermo) and mounted with ProLong Gold Antifade Mountant (Thermo). Imaging was done with an automated IX81 microscope (Olumpus) equipped with a MT20 illuminating unit with a 150 W mercury-xenon burner and a motorized stage (Merzhauser), a 20×/0.75 objective lens, and a 5-channel SEDAT filter set (Semrock). Data were acquired using ScanR Acquisition software (Olympus) and initial image analysis (segmentation, gating, intensity quantification) was done with ScanR Analysis (Olympus). Downstream analysis was done in R, using packages dplyr, tidyr, and data.table. Plots were generated with ggplot2. For each condition, 500 cells were analyzed.

For cell cytometry analysis, ER-tracker red (ER-Tracker™ Red, BODIPY™ TR Glibenclamide; for live-cell imaging, E34250, Thermo Fisher Scientific) was used according to the manufacturer’s instructions to selectively stain endoplasmic reticulum. Samples intensity were measured with BD LSRFortessa™ under FACS Diva Software v8.0.1 (BD) software control and analyzed using FlowJo (Data Analysis Software v10).

## Acknowledgments

We thank Ewa Borsuk for help with the generation of TENT5A and TENT5A-3xFLAG mouse lines, Katarzyna Prokop for assistance with cloning and all AD lab members for fruitful discussions and support. This work was supported by grant funding from Foundation for Polish Science (TEAM TECH CORE FACILITY/2017-4/5 to AD and TEAM TECH CORE FACILITY/2016-2/2), National Science Center (2019/33/B/NZ2/01773) to AD, as well as RVO 68378050 by Czech Academy of Sciences LM2015040 and LM2018126 for Czech Center of Phenogenomics provided by MEYS, CZ.02.1.01/0.0/0.0/16_013/0001789 Upgrade of the CCP: developing towards translation research by MEYS and ESIF, CZ.1.05/1.1.00/02.0109 provided by BIOCEV and MEYS and CZ.1.05/2.1.00/19.0395 Higher quality and capacity for transgenic models breeding by MEYS and ERDF to RS and JP.

## Authors contributions

OG analyzed the mouse phenotypes and performed all experiments on osteoblasts. GAN participated in mouse phenotyping and histological analysis, PK performed all bioinformatics analysis, SM performed DRS sequencing, tethering experiments, and supported the experimental design, MKK performed qRT-PCR and cytometry analyses, BT and AC performed microscopic analysis, FS and GAN performed microCT analysis, OB and OK performed SEM analysis, PS performed cryoEM analysis, DC performed mass spectrometry analysis, JG and MS constructed and genotyped mice, RS coordinated the mouse phenotyping pipeline at CCP, AD and JP conceived and directed the studies. OG and AD drafted the paper with the contribution of GAN and JP.

## Expanded View Figure Legends

**Expanded View Figure 1.** Selected phenotypes of TENT5A KO mice.

A–B. Genotyping of pups born from TENT5A(WT/-) × TENT5A(WT/-) heterozygotic matings on day 6 (A) and day 35 (B), performed for two independent cohorts.

C. Representation of wavy tail of the TENT5A KO mouse, which was present mainly in adult individuals.

D–E. Body weight analysis of WT, TENT5A heterozygotic, and TENT5A KO mice at 5 weeks old. (D) weight of males ((n=7–20); p < 0.0001, Unpaired t test with Welch’s correction). (E) weight of females ((n=6–19); p = 0.0043 for WT vs TENT5A KO; p < 0.0001 for TENT5A(WT/-) vs. TENT5A KO).

F. Alizarin Red/Alcian Blue staining of 8-week-old TENT5A KO skeleton with multiple ribs fractures visible.

G. Tibia and fibula dissected from WT and TENT5A KO adult mice. Bones were prepared by boiling and treatment with 30% hydrogen peroxide. Deformation of TENT5A KO tibia can be observed.

H. Alizarin Red/Alcian Blue staining of E18 embryo revealed no fractures. Five independent stainings were performed.

**Expanded View Figure 2.** Additional skeletal phenotypes

A. MicroCT image of a WT femur with selected regions of interest on metaphyseal trabecular bone (yellow) and mid-diaphyseal cortical bone (purple) for imaging and quantification. Scale bar, 1 mm.

B. MicroCT-derived femur morphometry results of trabecular bone total volume, trabecular thickness, and cortical thickness. Data are presented as mean ± SD. *P ≤ 0.05 (Student’s t-test).

C. Wider view of confocal scans of immunohistochemical staining for FLAG in femur sections from Tent5a-3xFLAG knock-in and control animals with FLAG in green and Hoechst in blue; insets denote parts of scans depicted in the main figure, Cx - cortical bone, BM - bone marrow; scale bar indicates 200μm;

**Expanded View Figure 3.** Additional TENT5A activity and substrate data

A. High-resolution northern blot analysis of SSR4 transcripts from SKMM1 cells transduced with TENT5AWT-GFP up to 72 h reveals that the SSR4 transcript is extensively polyadenylated by TENT5A.

B. Poly(A) tails added to reporter mRNA can be removed by RNase H treatment in the presence of oligo(dT)25. High-resolution northern blot analysis of RL mRNA from control HEK293 cells (lanes 1–2), after tethering of NHA-TENT5AaWT (lanes 3–4) or NHA-TENT5Amut (lanes 5 6).

C–D. DRS-based poly(A) lengths profiling of Rplp1 (C) and mtRnr2 (D) mRNAs from WT and TENT5A KO neonatal calvarial osteoblasts on day 14 of the maturation assay showing no significant changes in poly(A) lengths.

C. Rplp1: median poly(A) tail lengths (WT=66 nucleotides; KO=65 nucleotides).

D. mt-Rnr2: median poly(A) tail lengths (WT=9 nucleotides; KO=10 nucleotides).

E. Western blot analysis of SerpinF1 levels in mouse serum showing higher levels in WT than in TENT5A KO. Ponceau S staining was used as loading control.

F. Western blot analysis showing that expression of SPARC is higher in WT than TENT5A KO both on days 14 and 21 of the osteoblast maturation assay.

**Expanded View Figure 4.** TENT5A substrates are targeted to the ER and possesses relatively long poly(A) tails.

A. RT-qPCR analysis of TENT5C expression during the osteoblast maturation assay showing upregulation of TENT5C expression on days 7–21, normalized to the expression of HMBS (n=3–6; Mann–Whitney test: D0 vs. D7, p = 0.0159; D0 vs. D14, p = 0.0095; D0 vs. D21, p = 0.0159; D0 vs. D28, p = 0.0571).

B. Functional annotations (GO terms) significantly enriched for genes with short or long tails.

C: Fractionation of TENT5A-3xFLAG and WT osteoblasts followed by western blot analysis using anti-FLAG antibody. Cyt, cytoplasmic fraction; Mem, membrane fraction; Nuc, nuclear fraction.

D: 3’UTR motif analysis did not reveal any enriched motifs except for the canonical polyadenylation signal.

E: Substrates of TENT5A are characterized by higher than usual GC content.

F. Substrates of TENT5A are relatively short.

**Expanded View Figure 5.** Poly(A) tails distribution and TENT5A substrates in osteoblasts derived from long adult bones are similar to those of neonatals on day 14 of the maturation assay.

A. DRC based profiling of global poly(A) distribution in WT and TENT5A osteoblasts derived from adult long bones. WT median, 81 nucleotides; TENT5A KO median, 76 nucleotides.

B-C. DRC-based poly(A) lengths profiling of Col1a1 and Col1a2 mRNAs isolated from WT and TENT5A KO osteoblasts isolated from adult long bones.

B: Col1a1: median lengths of poly(A) tails (WT, 130 nucleotides; TENT5A KO, 98 nucleotides; p < 0.001).

C: Col1a2: median lengths of poly(A) tails (WT, 127 nucleotides; TENT5A KO, 108 nucleotides; p = 0.07).

D: Distribution of poly(A) tail lengths of mRNAs encoding extracellular matrix proteins (top) and other proteins (bottom) in WT and TENT5A KO adult long bones osteoblasts showing that mRNA encoding extracellular matrix proteins are the main targets of TENT5A. Extracellular matrix mRNAs medium poly(A) tail lengths (WT, 101; TENT5A KO, 78; other mRNAs: WT, 75; TENT5A KO, 64).

**Expanded View Table 1:** Analysis of differences in the lengths of poly(A) tails between WT TENT5A KO at D14

**Expanded View Table 2:** Differential expression analysis of TENT5A WT and TENT5A KO at D0 and D14.

**Expanded View Movie 1:** Video of 3D whole body scan of WT mouse rotating at the longitudinal axis

**Expanded View Movie 2:** Video of 3D whole body scan of TENT5A KO mouse rotating at the longitudinal axis where skeleton malformations are visible.

**Expanded View Movie 3:** Micro-CT-derived video showing longitudinal crossing through the WT trabecular bone. First, we observe the trabecular volume in yellow and then the trabecular spacing in a color heatmap.

**Expanded View Movie 4:** Micro-CT-derived video showing longitudinal crossing through the TENT5A KO trabecular bone. First, we observe the volume difference in yellow and then the trabecular spacing difference of TENT5A KO mouse.

**Appendix Table 1:** List of oligos used in this study.

**Appendix Table 2:** List of antibodies used in this study.

**Appendix Table 3:** Summary of DRS runs

